# Tau amyloid polymorphism is shaped by local structural propensities of its protein sequence

**DOI:** 10.1101/2022.10.24.512987

**Authors:** Nikolaos Louros, Martin Wilkinson, Grigoria Tsaka, Meine Ramakers, Chiara Morelli, Teresa Garcia, Rodrigo U. Gallardo, Sam D’Haeyer, Vera Goossens, Dominique Audenaert, Dietmar Rudolf Thal, Neil A. Ranson, Sheena E. Radford, Frederic Rousseau, Joost Schymkowitz

## Abstract

Different tauopathies are characterized by specific amyloid filament folds that are conserved between patients. Disease-specific tau filament folds probably reflect the specific pathological contexts leading to their formation including isoforms or post-translational modifications. Little is known, however, as to whether and how intrinsic conformational tendencies of the tau sequence itself contribute to its polymorphism. Using cryo-EM structure determination we find that a short amyloidogenic C-terminal peptide consisting of residues 350-362 of the tau repeat domain adopts the same polymorphic conformations in isolation as it does in the context of major disease-associated protofilament folds. Biophysical characterisation and molecular modelling show that the amyloid conformations adopted by this peptide constitute core structural motifs stabilizing distinct disease-associated tau filament folds. In accordance this segment also contributes to the efficient propagation of human AD tau seeds in tau reporter cells while it is irrelevant to heparin-induced recombinant seeds. Our findings suggest that tau 350-362 is key to the propagation of disease-associated tau polymorphs and that the conformational preferences of this segment predispose to the topological diversity observed in tau filament folds.

## Introduction

The intracellular deposition of tau in a non-native amyloid form characterizes a heterogeneous group of more than 20 neurodegenerative diseases called tauopathies including Alzheimer’s disease (AD), progressive supranuclear palsy (PSP) or corticobasal degeneration (CBD) among others^1^. Tau is most abundant in neuronal axons where it binds to microtubules, thereby contributing to microtubule assembly which is essential for the structural stability of neurons and axonal transport^2^. Under physiological conditions, tau is a monomeric, soluble, intrinsically disordered protein^3^. Pathological conditions lead to hyperphosphorylation and other post-translational modifications of tau, resulting in loss of microtubule binding and the aggregation of tau into intracellular amyloid deposits^4,5^. Once formed, tau amyloids act as seeds that facilitate the aggregation of additional tau monomers^6,7^. In addition, tau amyloid seeds can spread to functionally connected neurons where they perpetuate tau amyloid aggregation^6^. The exact role of tau amyloid deposition and spreading in neuronal cell death remains unclear. It is currently suspected that tau amyloids or their precursors aberrantly interact with cellular components including lipids, nucleic acids and proteins, leading to their functional dysregulation^8,9^. Tau amyloid deposits may also sequester other cellular components leading to neuronal impairment and cell death^8^.

Different tauopathies originate in different brain regions and neuronal cell types, resulting in distinct spatio-temporal disease progressions and neuropathological symptoms^10^, suggesting that the pathological mechanisms underlying neurodegeneration may vary between tauopathies. Recent high-resolution cryo-electron microscopy (cryo-EM) structures show that tau amyloids adopt different tertiary folds in different tauopathies, that are conserved between patients with the same pathology^11^. The correlation between neuropathology and polymorphism therefore suggests that polymorphism reflects disease-specific pathological events^12^.

Cellular conditions modifying tau polymorphism include post-translational modifications and alternative splicing^13^. However, it is unclear whether and to what degree the tau protein sequence itself predisposes to known disease polymorphs. Yet, this is key to understanding how amyloid structure responds to environmental perturbations including not only upstream biochemical events driving polymorph selection but also whether and how polymorphism determines disease-specific mechanisms of toxicity, selective cellular vulnerability and spatio-temporal patterns of spreading in the brain.

Tau consists of an N-terminal projection domain and a microtubule binding domain. The microtubule binding domain constitutes the core of tau amyloid deposits, while the N-terminal domain remains unstructured forming a ‘fuzzy coat’ around the amyloid core structure^14^. The microtubule assembly domain is composed of a tau repeat domain (tauRD with repeats R1 to R4), while alternative splicing results in isoforms with and without R2^15^. Two amyloid nucleating hexapeptide segments PHF6*(VQIINK_275-280_) in R2 and PHF6 (VQIVYK_306-311_) in R3 have been previously identified to drive amyloid assembly, as aggregation-suppressing mutations in these sequence segments strongly inhibit tau aggregation^16,17^. However, structural analysis revealed these segments adopt very similar conformations across polymorphs, suggesting that on their own they do not determine polymorphism^18-21^. At the same time, no amyloid-driving sequence has been identified yet in R4, despite this repeat is consistently present in all human disease tau polymorphs.

Here we identify a new amyloidogenic segment, PAM4 (for Polymorphic AMyloid segment in repeat 4), encompassing residues 350-362 of R4. Using tau aggregation reporter cells, we show that PAM4 induces aggregation of the tauRD domain. Cryo-EM shows that PAM4 fibrils are polymorphic, and that the conformations adopted by this segment in isolation are similar to the conformations PAM4 adopts in polymorphs from different human tauopathies. The conformations adopted by PAM4 in isolation are also thermodynamically favourable in the context of patient-derived amyloid structures, indicating that the tauRD domain possess intrinsic structural propensities favouring tau disease polymorphs. These findings, therefore, suggest that post-translational modifications and changes in splicing patterns reinforce rather than determine polymorphic preferences. This also suggests the initiation of amyloid aggregation will be extremely sensitive to such changes and that subtle cellular imbalances could be sufficient to direct polymorphism.

## Results

### The PAM4 fragment forms amyloid fibrils that seed aggregation in tau biosensor lines

The sequence-based amyloid prediction algorithm WALTZ^22^ identifies two previously validated aggregation prone regions in tauRD, namely the VQIINK^275-280^ and VQIVYK_306-311_ stretches known as PHF6* and PHF6, respectively^16,17^. However, WALTZ (**Fig. 1a, cyan area**) also predicts an additional putative aggregation motif spanning VQSKIGSLDNITH_350-362_ in R4 of tauRD domain which has not been previously characterized (hereafter referred to as PAM4 for **P**olymorphic **AM**yloid of repeat **4**). Given the ubiquitous presence of PAM4 in the structural amyloid cores of human tauopathies, we synthesized a peptide corresponding to this segment in order to experimentally determine its amyloidogenic properties. Thioflavin-T (Th-T) kinetics revealed that the PAM4 peptide forms aggregates in a concentration-dependent fashion (**Fig. 1b**). Thin films containing end-state aggregates were positively stained with the Congo red dye, as seen under bright-field illumination, and also exhibited a green birefringence that is typical for amyloid deposits when viewed under polarized light (**Fig. 1c**). Diffraction patterns produced from oriented fibrils of the peptide were indicative of a typical cross-β architecture, with a strong and meridionally oriented reflection at 4.7 Å and an equatorial reflection at 10.9 Å corresponding to the stacking and packing distances of β-strands and β-sheets along the fibril axis, respectively (**Fig. 1d**). In agreement, FTIR spectroscopy revealed two prominent peaks at 1631 cm^-1^ and 1680 cm^-1^ within the amide I region, both indicative of a parallel β-sheet-rich conformation (**Fig. 1e**). Morphological characterisation of fibrils formed by the PAM4 peptide, using atomic force microscopy (AFM) and transmission electron microscopy (TEM), revealed the formation of long and highly polymorphic amyloid fibrils (**Fig. 1f-g**). More specifically, in TEM we could identify multiple morphologies from straight and laterally interacting fibrils to twisted helical and ribbon-like morphologies, respectively (**Fig. 1f**). Similar morphologies were identified with AFM, in addition to certain hyper-twisted fibrils that were not picked up by TEM (**Fig. 1g, arrows**). We also investigated the amyloid nucleating properties of the same segment against the repeat domain of tau within a cellular context. To do this, we generated seeds by sonicating end-state PAM4 peptide amyloid fibrils and subsequently treating HEK-293 cells transiently expressing the tauRD conjugated to YFP. By counting the number of expressing cells showing a punctuate morphology through automated image analysis, the derived dose-response curve revealed a seeding efficiency of the PAM4 peptide (with a calculated EC_50_ = 4.1 μM) (**Fig. 1h-i**), showcasing its potency as a nucleating region of the tau protein.

**Figure 1.**
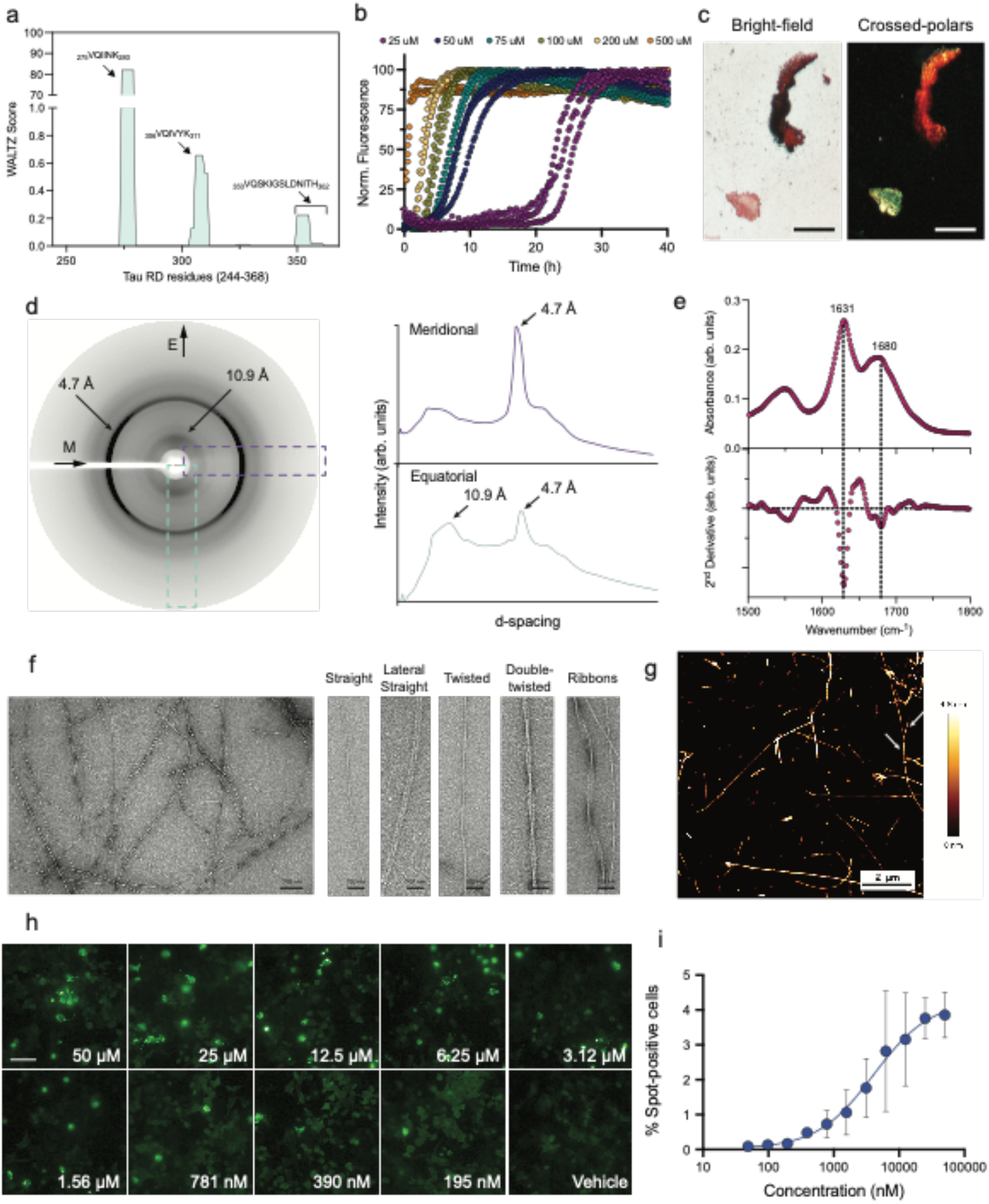
Biophysical characterisation of the amyloid-like properties of PAM4. (a) WALTZ aggregation propensity prediction for the tauRD. (b) Concentration-dependent Th-T kinetic assays of the PAM4 peptide (n=3 independent repeats). (c) Polarised microscopy reveals an apple-green birefringence shown by PAM4 peptide deposits which typically signifies the presence of amyloid aggregates. (d) Cross-β diffraction pattern produced by oriented fibres containing PAM4 peptide fibrils. Intensity quantification along the meridional (purple curve) and equatorial axis (cyan curve) of the pattern indicate the presence of an intense 4.7 Å and 10.9 Å reflection, respectively. (e) FTIR spectrum produced from fibril deposits of the PAM4 peptide. The prominent 1631 cm^-1^ and 1680cm^-1^ peaks are indicative of a dominant parallel β-sheet secondary structure. (f) Electron micrograph of fibrils formed by assembly of the PAM4 peptide. Higher magnifications of individual fibrils showcase the presence of highly polymorphic amyloid fibrils. (g) Atomic force microscopy image of PAM4 fibrils. Multiple helical morphologies can be observed in a single field of view. (h) PAM4 peptide seeds induce cellular seeding of a tauRD-YFP conjugate construct. (i) Concentration-dependent seeding quantification performed by counting the percentage of expressing cells containing fluorescent puncta (n=3 independent repeats).

### Structural determination by cryo-EM reveals the PAM4 fragment adopts polymorphic protofilament folds

We utilized cryo-electron microscopy for the structural determination of amyloid fibrils formed by the PAM4 peptide. First, we validated the highly polymorphic nature of the derived amyloid fibrils after performing two-dimensional (2D) class averaging and classification. This process revealed multiple distinct fibril polymorphs within the data (**Supplementary Fig. 1a**) with at least four major morphologies immediately identifiable (**Supplementary Fig. 1b**). The diversity in fibril architectures for the different PAM4 morphologies was striking. For further classification the dataset could be divided into two groups: one with the relatively simple twisting morphologies I and II and the second containing the wider, interlacing fibril segments of morphologies III and IV, which displayed more complex internal twisting structural elements (**Supplementary Fig. 1b and 1c**).

From the first group, only a single subtype corresponding to morphology I (representing 49% of the sub-population) was possible to resolve (**Fig. 2a-c and Supplementary Fig. 2**). Our final map, obtained at a resolution of 3.4 Å, reveals that this polymorph has a helical twist of 179.31° and a rise of 2.41 Å (**Fig. 2b**). The polymorph is composed of two identical protofilaments related by a pseudo-2_1_ screw symmetry axis (**Fig. 2a-c**). The second group was broadly characterised by a much wider average helical twist of 359° and an interlayer spacing of 4.82 Å, relating to the hydrogen bond distance between in-register β-sheets (**Fig. 2b**). From this, a further two PAM4 fibril structures were solved for the structurally related four-protofilament morphologies III and IV, at resolutions of 3.5 Å and 3.8 Å respectively (**Fig. 2a-c, Supplementary Fig. 2**) (**Table 1**). During the advanced stages of processing, a further three novel polymorphs were observed within the wider fibril group but could not be resolved to high resolution due to ambiguous helical layer separation within the maps (**Supplementary Fig. 2b**). Nonetheless, their respective protofilament cores clearly indicate peptide backbone conformations matching those determined within the solved morphologies III and IV with variations arising in the assembly of these related protofilaments within the context of the different fibril forms.

**Table 1.** Cryo-EM data collection, refinement and validation statistics for the PAM4 dataset.

|  | Morphology I<br>(EMDB-15550)<br>(PDB 8AOR) | Morphology III<br>(EMDB-15551)<br>(PDB 8AOS) | Morphology IV<br>(EMDB-15552)<br>(PDB 8AOT) |
| --- | --- | --- | --- |
| **Data collection and processing** |  |  |  |
| Magnification | 130,000 |  |  |
| Voltage (kV) | 300 |  |  |
| Detector | K2 |  |  |
| Electron exposure (e<sup>-</sup>/Å<sup>2</sup>) | 56 |  |  |
| Exposure rate (e<sup>-</sup>/pixel/s) | 8.0 |  |  |
| Nominal defocus range (μm) | -1.5 to -2.7 |  |  |
| Pixel size (Å) | 1.065 |  |  |
| Movies collected | 3,202 |  |  |
| Initial particle images (no.) | 560,812 |  |  |
| Final particle images (no.) | 23,707 | 9,085 | 5,955 |
| Map resolution (Å) | 3.4 | 3.5 | 3.8 |
| FSC threshold | 0.143 | 0.143 | 0.143 |
| Map resolution range (Å) |  |  |  |
| Symmetry imposed | C1 | C1 | C1 |
| Helical parameters |  |  |  |
| Helical twist (°) | 179.306 | 358.99 | 358.84 |
| Helical rise (Å) | 2.41 | 4.83 | 4.83 |
| **Refinement** |  |  |  |
| Initial model used (PDB code) | - | 8AOR | 8AOS |
| Map sharpening *B* factor (Å<sup>2</sup>) | -92 | -74 | -77 |
| Model to map correlation | 0.78 | 0.80 | 0.78 |
| Model composition |  |  |  |
| Non-hydrogen atoms | 1,224 | 2,448 | 2,448 |
| Protein residues total | 156 | 312 | 312 |
| Chains per helical layer | 4 | 8 | 8 |
| Helical layers modelled | 3 | 3 | 3 |
| *B* factors (Å<sup>2</sup>) |  |  |  |
| Protein | 73.0 | 65.0 | 87.1 |
| R.m.s. deviations |  |  |  |
| Bond lengths (Å) | 0.011 | 0.010 | 0.011 |
| Bond angles (°) | 1.088 | 0.848 | 1.070 |
| Validation |  |  |  |
| MolProbity score | 1.8 | 1.3 | 1.6 |
| Clashscore | 9.0 | 5.5 | 10.9 |
| Poor rotamers (%) | 0.0 | 0.0 | 0.0 |
| Ramachandran plot |  |  |  |
| Favored (%) | 95.7 | 100.0 | 100.0 |
| Allowed (%) | 4.3 | 0.0 | 0.0 |
| Disallowed (%) | 0.0 | 0.0 | 0.0 |

**Figure 2.**
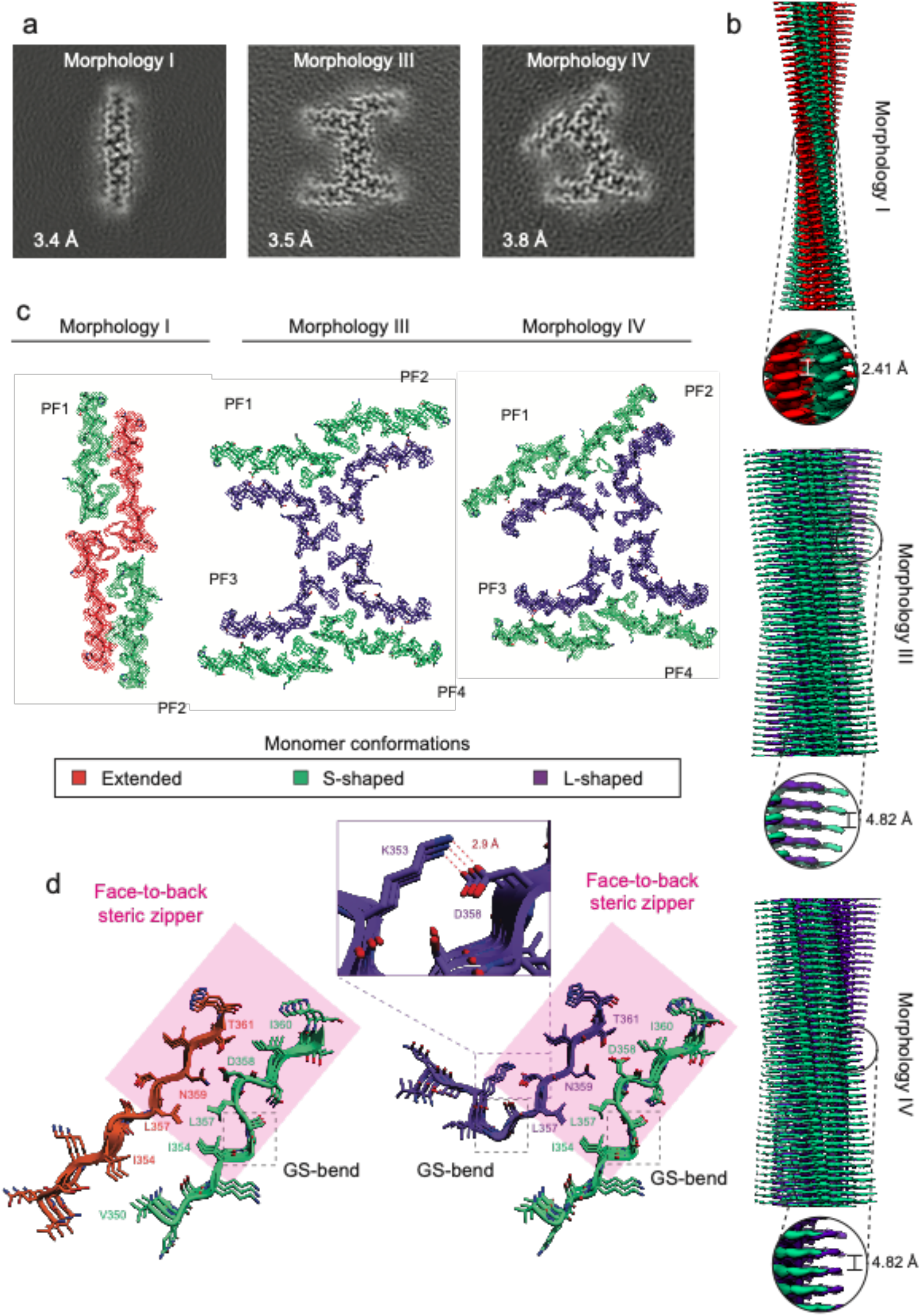
Structural determination of PAM4 fibrils using cryo-electron microscopy. (a) Projected slices of the different fibril morphology maps obtained after three-dimensional reconstruction. (b) Three-dimensional densities of the individual peptide polymorphs. (c) Cross-section of the structural layout corresponding to each morphology. Thinner fibrils are formed by four peptide monomers in an extended (shown in orange) or an S-shape conformation (shown in green), whereas wider morphologies are composed of eight peptide chains found in the same S-shaped or an L-shaped conformation (shown in purple), respectively. (d) A steric zipper interface (shown in magenta planes), formed between intercalating Ile354, Leu357, Ile360 residues and hydrogen bonded Asp358, Asn359 and T361 side chains, is conserved in all peptide polymorphs that in turn, differ due to structural heterogeneity introduced by Gly355 and Ser356 (shown in dashed boxes). The L-shaped monomer is further stabilised by an intramolecular salt bridge formed between Lys353 and Asp358, shown in a zoomed-in panel.

Cross-sectional analysis of the derived polymorphic PAM4 structures (**Fig. 2c**) reveals that the two protofilaments of the thinner morphology I fibrils incorporate two peptide chains packing laterally to form a dry steric zipper, reminiscent of a face-to-back conformation (**Fig. 2d**). This is a heterogeneous interface between a peptide chain in a fully extended conformation (**Fig. 2c-d, orange conformers**) and a less extended S-shaped conformation (**Fig. 2c-d, green conformers**). The S-shaped chain contains a GS-bend in the middle section of the backbone at the linker-like Gly355-Ser356 region which flips the N-terminal residues of this chain relative to those in the extended chain. The two protofilaments assemble in a head-to-head fashion, forming an interaction interface shaped by the Val350 residues, accompanied by cross-interacting Gln351-Gln351 ladders and an undefined density next to the N-terminal Val350 that may be attributed to HFIP molecules derived from the solute. Type III and IV fibril polymorphs are each composed of four identical protofilaments that are shared between both architectures (**Fig. 2c**). Each protofilament consists of two peptide chains forming an identical face-to-back steric zipper interface to that observed in type I protofilaments. As in the latter, one of the monomers presents a conserved S-shaped conformation featuring the GS-bend (**Fig. 2d**), but in this case it faces a monomer in a different conformation to the extended form seen in morphology I. Instead, resides a L-shaped monomer displaying a sharp angle turn at the same GS position to rotate the N-terminal residues ∼90° away from its interacting partner (**Fig. 2c-d, purple conformers**). This bent L-shaped conformation is further stabilised by a salt bridge formed between Lys353 and Asp358 residues on the surface of the protofilament (**Fig. 2d**). This facilitates a slightly altered head-to-head inter-protofilament packing relative to morphology I, which then repeats and assembles in two distinct orientations to generate the fibril morphologies III and IV respectively (further diversity in these relative orientations generate the fibril architectures visible within morphologies V and VII, **Supplementary Fig. 2b**).

### Polymorphs of the PAM4 peptide are representative of disease-associated tau polymorphs

Structural determination of the highly polymorphic PAM4 amyloid fibrils revealed that this segment can adopt three individual conformations in our preparations (**Fig. 3, E-, S- and L-conformations**) that assembles in different ways to generate multiple fibril morphologies (**Fig. 2b**). Both the Val350-Ile354 (**Fig. 3, N-term superposition**) and Leu357-His362 residue stretches (**Fig. 3, C-term superposition**) share overlapping dihedral angles between the three peptide monomers, which indicates that the intrinsic polymorphism of the fragment stems from the conformational instability introduced by the central Gly355 and Ser356 residues.

**Figure 3.**
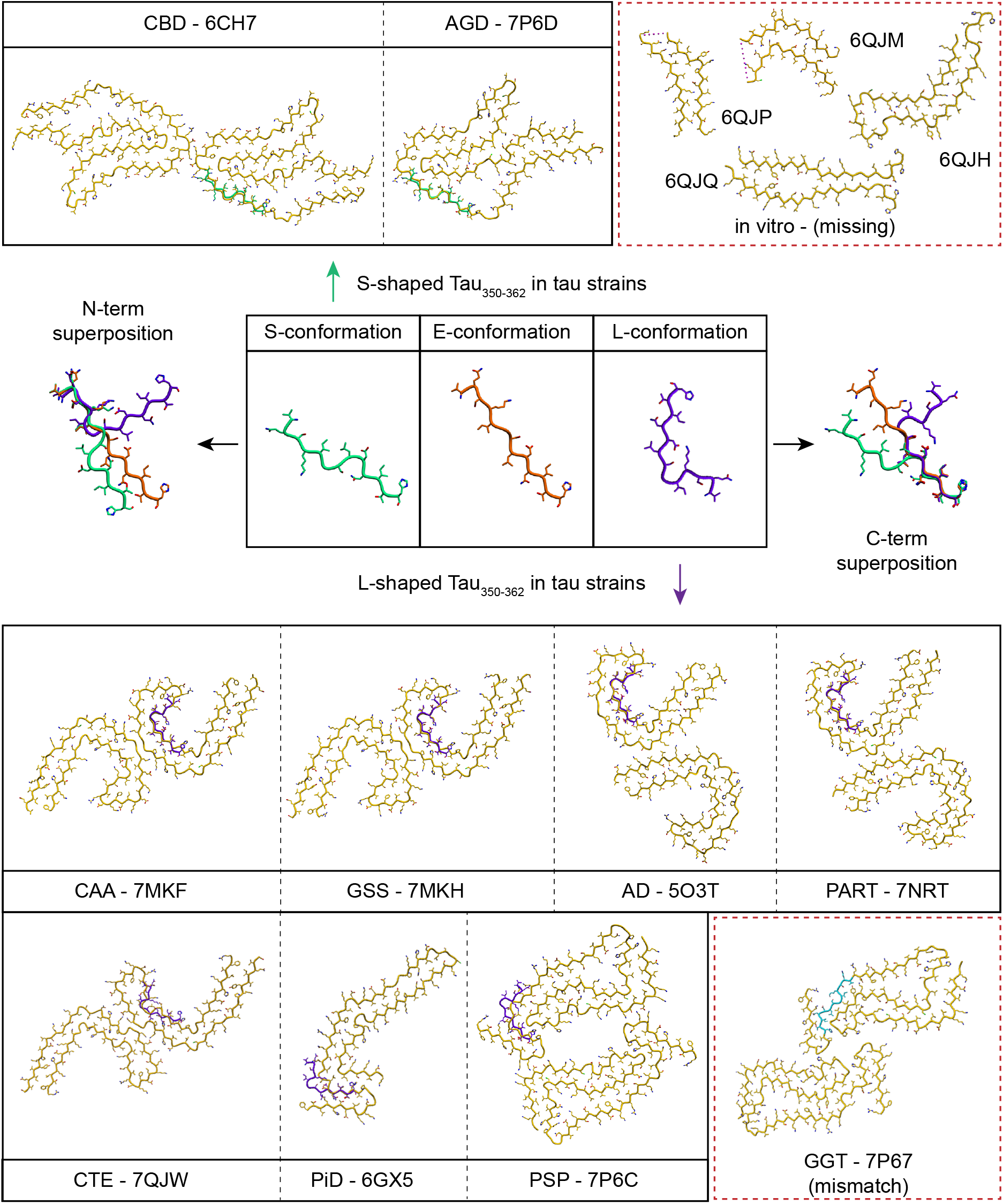
Polymorphism of ex vivo derived tau fibril aggregates traces back to the innate structural features of PAM4 fibrils. Three different monomeric conformation were identified for the PAM4 peptide in isolation, corresponding to an extended (E-conformation, shown in red), an S-shaped (S-conformation, shown in green) and an L-shaped (L-conformation, shown in purple) one. N- and C-terminal structural alignment highlight that the polymorphic instability of this segment stems from the central Gly355 and Ser356 residues. Top panel: Super-positioning of the S-shaped monomer to tau fibril structures isolated form CBD and AGD patients. In vitro tau strains that do not incorporate the segment in their fibril core are shown in the dashed box. Bottom panel: Structural alignment of the L-shaped conformation against tau fibril strains associated to CAA, GSS, AD, PART, CTE, PSP and Pick’s disease. No matching peptide conformer was identified in isolation for fibrils isolated for GGT patients, shown in a dashed red box. PSP conformers show structural differences to GGT conformers at the protofilament level only within the PAM4 region (shown in cyan), suggesting the importance of this segment in differentiating distinct disease-related tau polymorphs.

Superimposition of the PAM4 amyloid peptide conformations onto the structures of full-length human-derived tau amyloids reveals that the structural polymorphism of the PAM4 motif in isolation corresponds to its respective conformation within the protofilaments of major classes of human tauopathies including R3:R4, R1:R3:R4 and R2:R3:R4 tau isoforms. Specifically, the L-shaped conformation of the PAM4 peptide is retained in polymorphs linked to Alzheimer’s disease (AD), Gerstmann-Sträussler-Scheinker (GSS), primary age-related tauopathy (PART), cerebral amyloid angiopathy (CAA) and chronic traumatic encephalopathy (CTE), while it adopts similar L-shape angled conformations in progressive supranuclear palsy (PSP) and Pick’s disease (PiD) (**Fig. 3, bottom**). Additionally, the identified S-shaped PAM4 conformation matches to tau strains associated to corticobasal degeneration (CBD) and argyrophilic grain disease (AGD) (**Fig. 3, top**). In fact, in only one of the solved disease-associated tau amyloid structures to date, presented in globular glial tauopathy (GGT)-extracted strains, does the PAM4 motif adopt a novel conformation that was not observed in isolation (**Fig. 3, shown in cyan**). Of note, however, GGT protofilaments share structural similarity across the length of their core compared to PSP-derived conformers and differentiate primarily at a local interface shaped by the PAM4 region, as previously reported^12^. This differentiation stems from the Gly355-Ser356 stretch and highlights once more that this segment is crucial for distinguishing these polymorphs that are related to different tauopathies. The conserved polymorphic profile of PAM4, both as part of human-derived tau fibril cores and in isolation indicates that the structural nature of this fragment is intrinsic to its sequence and that polymorphic bias can be guided by context.

### Deletion of PAM4 impairs the cellular propagation of AD-patient tau seeds

Previous structural studies have shown that R4 of tau RD (which includes PAM4) is an integral part of AD-related tau protofilaments^23^ whereas it is absent from the ordered cores of heparin-induced recombinant fibrils^24^. We next asked whether PAM4 is relevant for the propagation of human AD-derived tau seeds. We therefore investigated the role of PAM4, but also PHF6 and PHF6*, on the propagation in cells of tau seeds derived from heparin-induced recombinant tau fibrils (rTau), as well as tau aggregates extracted from the brain of patients (3 individual cases) diagnosed with AD. To do so, we transiently expressed YFP-tauRD conjugates (incorporating the P301S mutation) missing either Tau_275-280_ (ΔPHF6*), Tau_306-311_ (ΔPHF6) or PAM4 (ΔPAM4) motifs in HEK293 cells and compared their induced intracellular aggregation to the full length tauRD(P301S) construct, using both sources of tau seeds (**Fig. 4a**).

**Figure 4.**
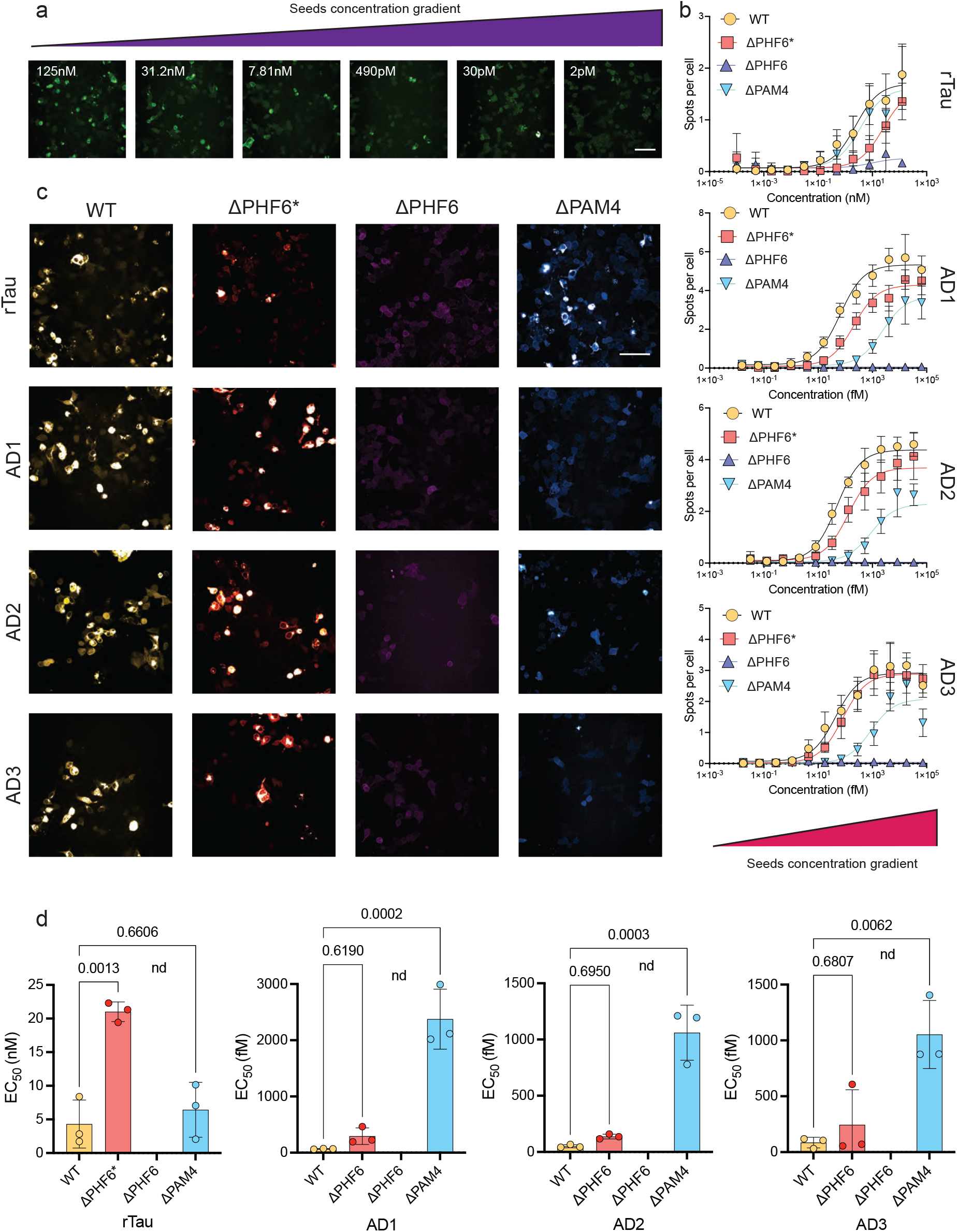
Cellular screening using deletion constructs of the tauRD reveals a differential relationship of APRs to tau aggregate strains. (a) Intracellular seeding of cells expressing tauRD-YFP is reported by counting the number of cells with a punctuate morphology in a concentration dependent manner upon treatment with rTau seeds. (b) Dose-response curves after the treatment of cells expressing tauRD (WT), ΔPHF6*, ΔPHF6 or ΔPAM4 with various concentrations of rTau or extracts isolated from three independent AD cases. (c) Representative images of treated cells with selected seed concentrations (shown in arrow in (b)). (d) Inverted effects of the ΔPAM4 and ΔPHF6* on seeding efficiencies, shown as changes in EC_50_ values, of recombinantly produced or AD extracted tau aggregates validates indicate that a bias towards specific tau polymorphs, in contrast to ΔPHF6 that is generally critical for tau aggregation. Bar plots represent mean values ± SD (n=3 independent samples).

Dose-response curves of seeding efficiency, reported as the number of expressing cells containing fluorescent puncta, showed that Tau seed propagation was completely abolished in ΔPHF6 expressing cells, regardless of the source of the seed aggregates (**Fig. 4b-c, purple curves**). This finding corroborates the crucial role of PHF6 in tau amyloid aggregation both in vitro and in vivo^16,17^. Also, in agreement with previous structural cryo-EM studies, we found the PHF6* motif was contributing to the propagation of *in vitro* generated tau seeds but not the propagation of seeds from AD patients. Specifically, a significant reduction in seeding efficiency was observed in ΔPHF6*-expressing cells when treated with rTau seeds (**Fig. 4b-c, red curves**), resulting in a 5-fold decrease of the EC_50_ value of ΔPHF6* cells when compared to full length tauRD(P301S) (**Fig. 4d**). Conversely, no significant changes in seeding efficiency were observed when treating cells expressing the ΔPAM4 construct with heparin-prepared seeds (**Fig. 4c-d**), showing that PAM4 does not contribute to the propagation of heparin-induced recombinant tau polymorphs.

Whereas the EC_50_ of rTau seed propagation in our assay lie in the nM range, AD-patient seed propagation potential was much higher displaying EC_50_ values in the fM range (**Fig. 4d**). Importantly, and in contrast to what was observed for rTau, seeding efficiency of AD extracts remained mostly unaffected upon deletion of PHF6*, pinpointing that it is not required to propagate AD polymorphs. On the other hand, deletion of PAM4 significantly impaired the ability of AD extracts to induce seeding in a cellular context. This translated to a more than thousand-fold difference in seeding efficiencies for each of the three individual cases tested in our assay, measured as the comparison of EC_50_ values between cells expressing the unmodified repeat domain of tau and those missing the corresponding PAM4 motif (**Fig. 4d**). In conclusion, our cellular findings confirm that not only structurally, but also kinetically, heparin-induced rTau amyloids are not representative of patient-derived AD tau amyloids. Importantly, they also demonstrate the key role of the R4 domain and in particular its PAM4 segment for the propagation of AD-patient tau seeds. Finally, our findings show that the cryo-EM amyloid core structures of these two tau polymorphs can be leveraged to understand the structure-activity relationship between amyloid core structure and its amyloid propagation in cells.

### PAM4 segment polymorphs contribute to the stability of disease-associated tau polymorphs

The PAM4 peptide in isolation populates specific structural amyloid folds, implying that these are thermodynamically the most stable conformations for the PAM4 sequence. These conformations are similar to those that the PAM4 segment adopts in the amyloid core of disease-associated tau polymorphs. The question then is whether these intrinsic conformational preferences of PAM4 remain thermodynamically stabilizing in the context of full-length amyloid cores and thus whether the structural preferences of PAM4 contribute to tau polymorphism. Using Stamp-DB, a public repository for amyloid fibril polymorph classification, we extracted and analysed the distribution of per residue energetic contributions along the fibril cores of all experimentally determined tau polymorph structures^25^. This repository utilises the FoldX force field for thermodynamic calculations^18,26^, which does not consider potential post-translational modifications, nor does it consider possible stabilizing interactions by unresolved non-proteinaceous electron densities in these amyloid structures. The energy profiles generated by FoldX, therefore, report only on how intrinsic polypeptide main-chain and side-chain interactions (i.e. cross-β quaternary structure and the way in which these are stabilized by tertiary interactions) contribute to the stability of the amyloid core.

A cumulative profile of tau fibril energetics displays the PAM4 motif as a significantly stabilising region of the sequence within the fibril structures (**Fig. 5a**). Additionally, it indicates that beyond PHF6, PHF6* and PAM4, other segments of tauRD also – to a lesser degree - contribute to the stability of different protofilament folds. These findings are correlated by CORDAX, a structure-based machine learning predictor that relies on cross-β compatibility^27^, which predicts two of these novel segments comprising residues 326 to 331 (sequence GNIHHK) and 344 to 349 (sequence LDFKDR) as well identifying part of the PAM4 motif (sequence LDNITH_357-362_) (**Fig. 5b, purple line**).

**Figure 5.**
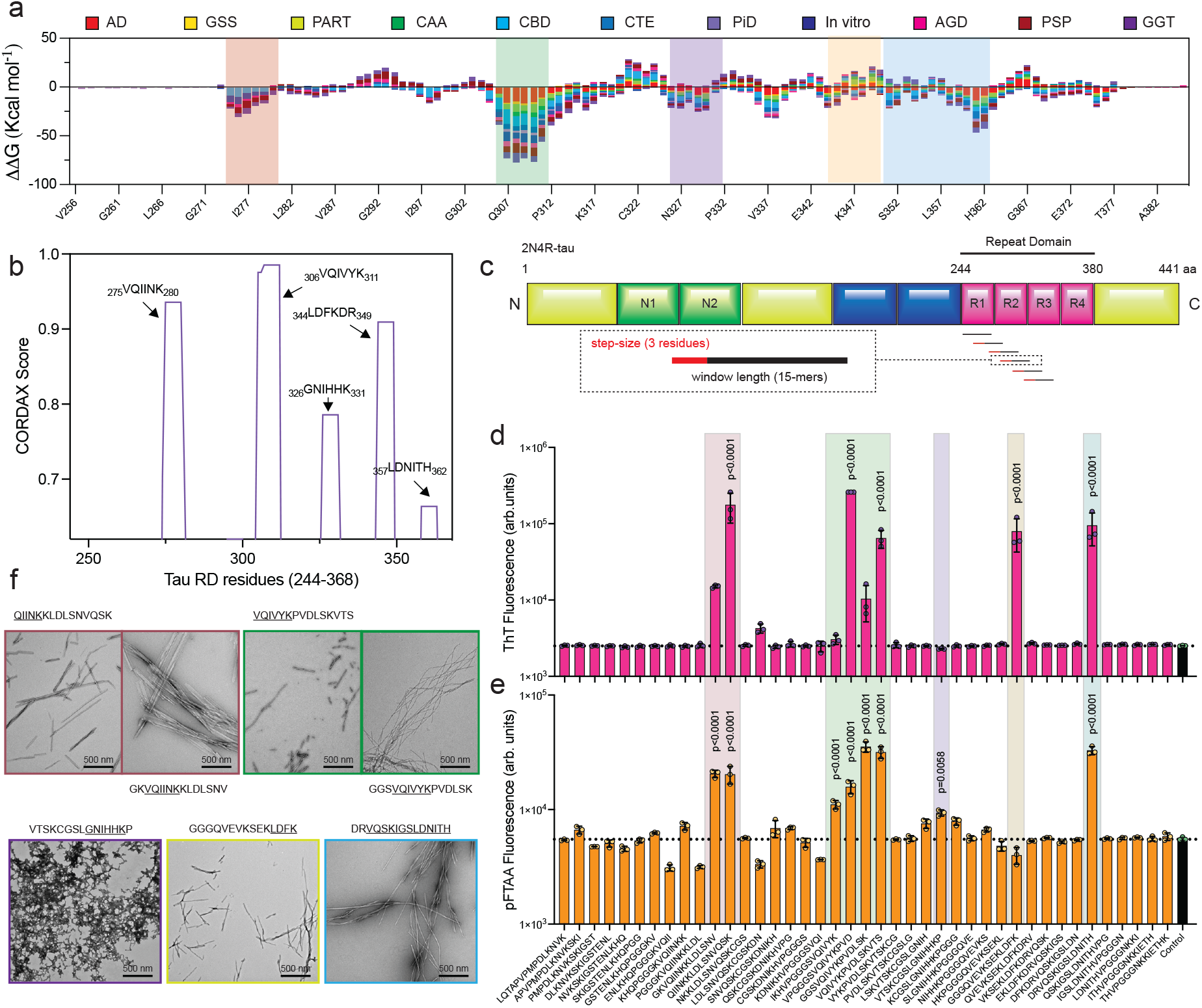
Profiling the cross-β compatibility of the tau repeat domain. (a) Stacked bar plot indicating the stacked individual stability contributions provided by each residue for tau fibrils polymorphs (colour-coded per disease). (b) Aggregation score profile of the tau repeat domain, as predicted by CORDAX (purple line). (c) Sliding window approach (15-residue windows incorporating 3-residue increments) used to generate a peptide library spanning the tau repeat domain. (d-e) End-state fluorescence analysis using Th-T (top) and pFTAA (bottom). Peptides showing increased fluorescence compared to the vehicle control (shown in black) were identified as positive for aggregation (windows containing each one of the predicted APRs are shown in colour-shaded areas). Bar plots indicate mean values +/- SD (n= 3 biologically independent samples). Statistical significance was determined using one-way ANOVA with Tukey’s test for multiple comparisons. (f) Electron micrographs validating the formation of amyloid-like fibril aggregates by the identified peptide windows. Colour-coded outlines matching the shaded-areas shown in (d-e) are used to highlight the corresponding predicted APRs contained in each peptide window.

In order to validate these predicted sequence profiles, we experimentally screened the amyloid propensity of the entire tauRD domain with a library of 41 peptides covering the entire tauRD region from N to C-terminus, using a sliding window strategy of 15-mer peptides with three-residue step-size increments (**Fig. 5c**). The amyloid propensity of each peptide was determined by end-state fluorescence with two different amyloid-reporting dyes. As electrostatic repulsions have been shown to often result in suboptimal amyloid-dye binding interactions^28^, we selected the positively charged Th-T and the negatively charged pFTAA, respectively. The amyloid propensity screen of tauRD segments (**Fig. 5d & e**) revealed amyloid structure in two adjacent peptides spanning residues 273-290 both containing PHF6*(red-shaded area), 4 adjacent peptides encompassing residues 297-320 all containing PHF6 (green shaded area) and one peptide spanning residues 348-362 containing PAM4 (cyan-shaded area). These three regions coincide with the most stable segments of our thermodynamic profile across polymorphs (**Fig. 6a**). Electron microscopy validated that those peptides containing either PHF6*, PHF6 or PAM4 form typical long and unbranched amyloid-like fibril aggregates (**Fig. 5f, green and red-boxes**). Of note, neighbouring windows containing partial segments of the PAM4 motif did not produce any type of aggregates, suggesting that the intact motif is required for the formation of stable amyloid aggregates in isolation.

**Figure 6.**
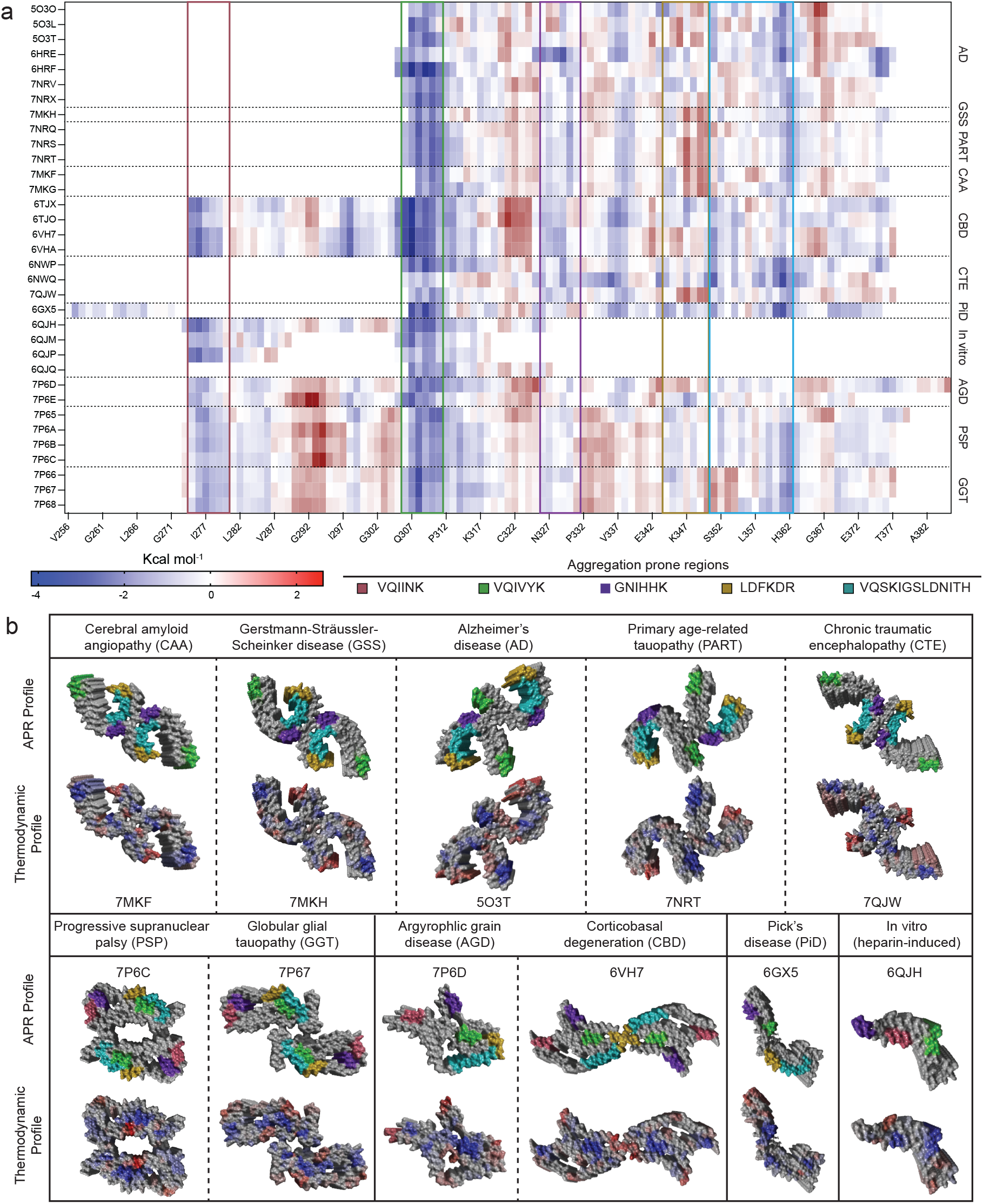
Analysis of the thermodynamic potential of tau fibril polymorphs reveals specific regions of stability. (a) Heatmap indicating the per residue energy contributions (shown in Kcal mol^-1^ as shown in the legend) for all structurally determined tau amyloid fibril cores. Left axis indicates individual structures listed with their respective PDB codes. Right axis includes a grouping of structures per pathology and the x-axis indicates tau residue numbers. Boxed outlines highlight the APR regions identified in this work, colour-coded as shown in the legend. (b) Surface representations of characteristic polymorphs per disease, colour-coded as shown in the legend to indicate either APR positioning (top) or per residue stability contribution (bottom), respectively.

Two additional amyloidogenic segments were also detected in the amyloid propensity fluorescence screen. The first segment consists of the peptide spanning residues 318-332 (**Fig. 5d & e**, purple-shaded area) containing the GNIHHK_326-331_ sequence predicted by CORDAX and highlighted by the FoldX/Stamp-DB thermodynamic profile to be stabilising across the tau fibril polymorphs. Electron microscopy showed that this peptide did not assemble into ordered amyloid-like filaments but rather formed amorphous aggregates (**Fig. 5f, purple box**) suggesting that while aggregation prone, this sequence needs additional interactions to form regular cross-β structure. The second additional amyloidogenic segment consists of the peptide spanning residues 333-347 (**Fig. 5d & e**, yellow-shaded area **box**), for which electron microscopy showed that it forms unbranched amyloids (**Fig. 5f, yellow box**). This peptide contains the previously predicted VEVKSE_337-342_ motif^29^, as well as the LDFKDR_344-349_ sequence predicted by CORDAX. Again, this agrees with our thermodynamic profile in which both regions constitute stable segments. Together, our data validate our thermodynamic calculations pinpointing regions of high stability in tau amyloid protofilament cores and that they generally reside along segments that possess local cross-β propensity as isolated peptides. Furthermore, this analysis shows that tau amyloid protofilament cores are not homogenously stable, but instead possess segments of high and low structural stability, as not all its constituent residues spontaneously adopt the amyloid conformation as single peptides.

A closer look at the per residues energy contributions confirmed that, in the context of full-length tau fibril cores and in line with previous reports^30^, PHF6 is the most stable sequence segment across all polymorphic structures, including in vitro and in vivo polymorphs. Furthermore, PHF6* also contributes to the stability of polymorphs with longer protofilament cores in which it is structurally incorporated including CBD, AGD, PSP, GGT, as well as in heparin-induced polymorphs for which our cellular data highlighted its importance (**Fig. 6**). While PHF6* and PHF6 are the most stable structural segments of R2 and R3, respectively, it also appears that PAM4 is the most stable segment of R4 in all polymorphs in which it is present (**Fig. 6a & b**). The fact that the local stability of PAM4 amyloids is not overruled by tertiary interactions in full-length amyloid cores suggests that the intrinsic structural propensity of PAM4 restricts the available conformational freedom of tau protofilament folds to those that are preferred by PAM4. The way in which the PAM4 conformations are incorporated in tau fibril polymorphs is in good agreement with the classification recently proposed by Shi et al^12^. R3:R4-containing polymorphs, including AD, PART and CTE protofilament folds, are stabilized by the L-shaped PAM4 polymorph (**Fig. 3 and Fig. 6b**). In particular, the L-shaped conformation of PAM4 stabilizes one leg of the horseshoe structure of these polymorphs, forming a stabilising heterotypic interface with GNIHHK_326-331_, while PHF6 stabilizes the other leg. The R1:R3:R4-containing structure of Pick’s disease also adopts a two legged - albeit less curved - protofilament fold, where again PHF6 stabilizes one leg, while PAM4 stabilizes the other (**Fig. 3 and Fig. 6b**). The S-shaped PAM4 polymorph is found in R2:R3:R4-containing polymorphs including AGD and CBD. These protofilament folds incorporate a longer section of the tauRD, but adopt a more compact structure, the core of which is stabilized by both PHF6 and PAM4 and whereby the N-terminal PHF6* closes the structure by interacting with the C-terminus of the amyloid core. Finally, PAM4 forms a heterotypic steric zipper interface of exceptional stability with PHF6 in PSP amyloid cores and is conversely stabilised by a similar interface in GGT polymorphs where it corresponds to poor PAM4 energetics. This may potentially explain why the novel PAM4 conformation seen in the GGT fibril structure was not recovered in isolation.

## Discussion

The atomic structure of tau inclusions found in patients of more than a dozen different tauopathies have been solved by cryoEM^11^. The major insight of these efforts has been that tau amyloid polymorphs are disease-specific and conserved between patients. It is still unclear what determines the variety of possible tau polymorphs and how specific polymorphs are selected in disease. Tau isoform composition appears to be the primary source of polymorphism^19,20,24,31-33^. Yet, tau inclusions from pathologies with similar isoform composition (e.g. CBD and PSP or AD and CTE) still adopt different filament folds indicating that other factors also shape the diversity of tau filament folds^20,33^. Disease-specific post-translational modifications^31^, as well as other physiopathological cellular interactions probably play an important role for the maturation of pathological tau into disease specific polymorphs^8,34^.

Here we find that - independent of isoform composition or post-translational modifications – local structural propensities of the tau sequence itself orients the amyloid structural diversity observed in human pathological tau inclusions. We identified PAM4, a short segment of consisting of residues 350-362 in tau repeat 4, that is able to switch between alternative amyloid conformations independently of the rest of the tau protein sequence. Indeed, despite its short length, a 13-mer peptide encoding the PAM4 sequence displays a surprising degree of conformational polymorphism. At the same time PAM4 polymorphism is not degenerate adopting a restricted number of at least four well-defined conformations. This suggests these conformations constitute relatively stable local structural motifs that are reminiscent of short supersecondary motifs, such as β-turns in globular proteins. Structural inspection of the PAM4 atomic structures shows how VQSKI and LDNITH provide stability to the cross-β assembly along the PAM4 filament axis while the flexibility of the GS linker provides for alternative backbone orientations perpendicular to the filament axis. Importantly, the local amyloid structural motifs adopted by the PAM4 peptide are maintained in the context of different human pathological tau filament folds. This suggests that PAM4 motifs constitute stable structural elements of mature disease polymorphs. Structural thermodynamic analysis of tau filament folds confirms that PAM4 motifs constitute core structural elements that both stabilize cross-beta assembly along the filament axis, as well as alternative tertiary packing of the filament folds. The importance of PAM4 for the local stability of various tau filament folds is further highlighted by the fact that local PAM4 polymorphism does not align with the hierarchical classification of tau filaments: the L-shaped PAM4 conformation, for example, is found in R3:R4 (AD, CTE and PART), R1:R3:R4 (Pick’s disease) and R2:R3:R4 (PSP) polymorphs, while both the L-shaped and S-shaped PAM4 polymorph are found in different subclasses of R2:R3:R4 tau filament folds (CBD vs AGD, respectively). The importance of PAM4 for the stability of tau inclusions is further supported by the finding of Eisenberg and co-workers that EGCG binding disassembles tau filaments by introducing a wedging effect that causes a backbone density breakage at Gly355, arguing that this could be an important starting step for tau fibril dissociation^35^. Moreover, Gly355 introduces a ∼70° turn that is recapitulated in the L-shaped monomer in isolation and has been shown to be critical for establishing the β-helical geometry of AD fibrils^20^. This β-helix structure has been linked to promoting the prion-like propensity of C-shaped tau strains^20^ and is further stabilised by a salt bridge between Lys353 and Asp358 stacked ladders^20,36^ in the L-shaped conformer. Phosphorylation of Ser356, on the other hand, has been shown to reduce in vitro fibrillation and cellular seeding of tau^37^, as well as change the dissociation free energy at the ends of AD filaments^38^, suggesting that the structural flexibility provided by this segment is important for tau fibril stabilization. Together, our findings suggest that the short PAM4 sequence is both amyloidogenic and polymorphic and that it is a key stabilizing element of tau pathological inclusions.

PAM4 is part of tau repeat R4 which, together with R3, is present in all known human pathological tau inclusions^12^, while it is absent of heparin-induced filaments^24^. We experimentally investigated the relationship between PAM4 and previously known amyloid-nucleation determining hexapeptides PHF6 and PHF6* of repeat R2 and R3, respectively using tau amyloid reporter cells. Strikingly, recombinant seeds have EC50 in the nM range while human patient extracted AD seeds have EC50 in the fM range. The origin of this huge gap in seeding activity remains to be further investigated, but it is in line with the hypothesis that human samples contain additional co-factors favouring tau seeding^39^. Nevertheless, deletion of PHF6 completely suppresses the ability of tau to be seeded by both heparin-induced as well as pathological AD tau seeds, confirming the dominant role of PHF6 for tau filament assembly^40^. While PAM4 deletion does not entirely abolish AD seeding, it still reduces seeding efficiency by three orders of magnitude. As expected from structure^20,24^, PHF6* plays no role in the seeding efficiency of AD patient extracted material, while conversely PAM4 has no relevance for the seeding efficiency of heparin-induced seeds. The lower impact of PAM4 on tau seeding efficiency probably results from the higher entropic cost of immobilizing PAM4 into a specific conformation. Indeed, while PAM4 acts as a conformational switch between its S and L conformation, the conformation of PHF6 is virtually identical in all polymorphs. This could potentially further be investigated by mutants that favour either conformation.

Our screening of the cross-beta propensity of the entire tau repeat domain revealed that other regions of the protein could act as PAM4 further contributing to the topological constraints shaping tau amyloid filament folds. Of notice, it was previously found that local conformational propensities around the PGGG spacers between tau repeats protect against tau amyloid assembly probably by forming turns that bury nucleating regions, such as PHF6 and PHF6*^30,41^. Tau is an intrinsically disordered protein the structural dynamics of which have been evolutionary selected for function. The complexity of the local structural propensities embedded in its sequence also contribute to the polymorphic nature of tau amyloid inclusions as well as to how tau responds to disease-specific contexts^21^. Understanding these reciprocities will be key to elucidating the role of polymorphism in human tauopathies.

## Methods

### Computational predictions

Calculations of aggregation propensity were performed using state-of-art aggregation predictors, such as WALTZ^22^ and Cordax^27^. WALTZ is based on a position-specific scoring matrix produced against a wide dataset of amyloidogenic sequences, whereas Cordax is a structure-based predictor that is less limited by typical propensities such as beta-propensity and hydrophobicity and as a result, is able to provide predictions that untether aggregation propensity from solubility. Thermodynamic profiles of amyloid fibril structures were calculated as described previously^18^ and were extracted from StAmP-DB, a structural database for polymorphic amyloid fibril structures^25^.

### Peptide library screening

The peptide library spanning the tau repeat domain residues was synthesized from GenScript, ensuring each peptide was of sufficient purity (>90%) and with blocked ends (N-terminal acetylation, C-terminal amidation). Samples were initially pre-treated with 1,1,1,3,3,3-hexafluoro-isopropanol (HFIP) (Merck), then dissolved in 25mM HEPES, 10mM KCl, 5mM MgCl_2_, 3mM TCEP, 0.01% NaN_3_, pH 7.2 buffer. A synthetic peptide corresponding to the PAM4 region was synthesised using an Intavis Multipep RSi solid phase peptide synthesis robot. Peptide purity (>90%) was validated with the use of RP-HPLC purification protocols. Peptide aliquots were stored as ether precipitates (−20 °C). After initial HFIP pre-treatment, samples were dissolved in milli-Q water.

### Negative staining

Hits derived from the peptide screen library were incubated for 7 days at room temperature. A drop (5 μL) of each peptide solution was added on glow-discharged (50s) 400-mesh carbon-coated copper grids (Agar Scientific Ltd., England). The grids were subsequently washed several times with milli-Q water and negatively stained using a 2% (w/v) uranyl acetate solution (in milli-Q). Grids were examined with a JEM-1400 120 kV transmission electron microscope (JEOL, Japan), operated at 80 keV.

For the preparation of grids for the PAM4 peptide, a 5 mg/mL peptide solution containing end state fibrils was diluted with milli-Q water before application to 300 mesh continuous carbon grids for 60s. The sample was blotted, washed with water and then stained with 1% (w/v) uranyl acetate. Grids were imaged using a Technai F20 microscope equipped with a FEI Ceta CMOS detector.

### Thioflavin-T and pFTAA binding assays

Solutions of each peptide (200 μM) were prepared and analysed in half-area black 96-well microplates (Corning, USA). For PAM4 kinetics, samples with various concentrations of the peptide were prepared in parallel, following filtering with 0.2 μΜ filters. Aggregation was assessed by adding Th-T (Sigma) or pFTAA (Ebba Biotech AB) at a final concentration of 25 μM and 0.75 μM, respectively. Fluorescence intensities were measured as independent repeats (n=3) at 30°C, with a double-orbital shaking step (100rpm for 10s) before each cycle, using a FLUOstar Omega plate reader (BMG Labtech, Germany), equipped with an excitation filter at 440 nm and corresponding emission filters at 490 and 510 nm, respectively.

### Congo red staining

Droplets (10 μL) of the PAM4 peptide solution containing mature amyloid fibrils were dried slowly in ambient conditions on glass slides in order to form thin films. The films were then stained with a Congo red (Sigma) solution (0.1% w/v) prepared in milli-Q water for 20 min. De-staining was performed with gradient ethanol solutions (70–90%). Imaging was performed on a SMZ800 stereomicroscope equipped with a polarizing filter (Nikon) and a DS-2Mv digital camera (Nikon).

### X-ray diffraction

A single droplet of the PAM4 peptide solution containing mature amyloid fibrils was dried between wax-covered capillary tubes in order to form oriented fibres suitable for X-ray fibre diffraction. X-ray diffraction patterns were collected with a Rigaku (Tokyo, Japan) copper rotating anode (RA-Micro7 HFM) operated at 40 kV and 30 mA, with a wavelength λ = 1.54 Å. The specimen-to-film distance was set to 200 mm and the exposure time was 1200 s. Diffraction patterns were studied using and displayed using the Adxv software (Scripps Research, USA).

### Fourier-Transform Infrared Spectroscopy (FTIR)

Suspensions (5 μL) of the PAM4 peptide were cast onto a 96-well silicon microplate (Bruker) and air-dried to form thin films. Spectral scans (120) were acquired at 4 nm^-1^ resolution in transmission mode and averaged to improved signal-to-noise ratio, using an HTS-XT FTIR microplate reader (Bruker). Background correction was performed by subtracting spectra obtained from a blank position of the microplate. Spectral normalisation and 2^nd^ derivatives with a 13-point smoothing, using Savitzky-Golay filtering^42^, were calculated using the OPUS software.

### Atomic Force Microscopy (AFM)

Solutions containing fibrils formed by the PAM4 peptide were diluted in milli-Q water to a final concentration of 1mg mL^-1^, then adsorbed to freshly cleaved mica for 15 min at room temperature. The mica was carefully washed 3 times (milli-Q water), to remove non-adsorbed material, and left to dry for 30 min at ambient conditions. For AFM scanning, RFESP-75 (with a normal frequency of 75kHz, a spring constant of 3 N/m and tip radius of 8nm) cantilever probes (Bruker) were used. Amyloid fibrils were imaged using a JPK NanoWizard Sense+ (Bruker) in oscillation mode (scan rate 0.5 Hz; amplitude setpoint 0.47 V), at a scan size of 10×10 um with an IGain of 300 Hz.

### Cryo-EM data collection

Fibrillated PAM4 peptide was diluted to 0.5 mg/mL in MilliQ water prior to application onto Tergeo plasma cleaned (Pie Scientific) Lacey carbon 300 mesh grids. The sample was blotted and frozen in liquid ethane using a Vitrobot Mark IV (FEI) with a 5 s wait and 5 s blot time respectively. The Vitrobot chamber was maintained at close to 100% humidity and 8°C. The cryo-EM dataset was collected using a Titan Krios electron microscope (Thermo Fisher) operated at 300 kV with a Gatan K2 detector in counting mode. A nominal magnification of 130,000x was set yielding a pixel size of 1.065 Å. A total of 3,202 movies were collected with a nominal defocus range of -1.5 to -2.7 µm using a total dose of ∼56 e-/Å^2^ over an exposure of 8 s, which corresponded to a dose rate of ∼8 e^-^/pixel/s.

### Cryo-EM data processing

Each 32-frame movie stack was aligned and summed using motion correction in RELION3^43^ and CTF parameters were estimated for each micrograph using CTFFIND v4.14^44^. Poor quality images, based on CTF figure of merit, defocus value and predicted resolution were removed to give 2,312 micrographs for further processing. Fibrils from 93 micrographs were manually picked in RELION and the extracted segments used to train automated filament picking in crYOLO^45^. Using an inter-box spacing of four helical repeats (∼19 Å), a total of 560,812 helical segments were extracted 2x binned in RELION with box dimensions of ∼850 Å. 2D classification was used to separate out picking artefacts and unfeatured objects to leave 483,274 fibril segments. During the next round of 2D classification, the class averages were split into 2x subset pools based on the apparent fibril physiology, with 249,716 segments saved as simple twisting fibrils (morphology I & II) and 226,088 segments saved with more complex interlacing structural elements (Morphology III & IV) (**Supplementary Fig. 1**). Each separate pool was cleaned by 2D classifying once more 2x binned and then a final time after unbinned extraction with 320 Å box dimensions to remove unfeatured and mis-classified fibril segments.

The cleaned 171,497 type I and Type II fibril segments were used for 3D classification using a 20 Å low-pass filtered initial template generated from a representative 2D class average with an initial helical twist and rise of 358.7° and 4.81 Å respectively (**Supplementary Fig. 2**). From this, a single class containing 84,939 segments (type I, 49% of those classified) was selected containing a resolvable fibril core structure. Initial 3D refinements confirmed that this polymorph contained a pseudo-2_1_ screw symmetry axis before two more rounds of 3D classification filtered the subset down to 26,012 ordered segments. After multiple sequential 3D refinements with searching of the helical parameters and CTF refinement of the per-particle defocus estimates, a final type I morphology map was obtained at a resolution of 3.4 Å (RELION gold-standard with 0.143 FSC cutoff) with a helical twist of 179.31° and rise of 2.41 Å and sharpening B-factor of -92 Å^2^. The structure of the type II segments that additionally appeared in the first 3D classification run could not be resolved despite thorough classification and searching of the helical parameters both with and without a screw axis.

The cleaned 165,342 Form2 and Form3 segments were used for 3D classification using a 15 Å low-pass filtered initial template generated from a representative 2D class average with an initial helical twist and rise of 359.0° and 4.82 Å respectively (**Supplementary Fig. 2**). The resulting classes could be sub-divided further to select 30,585 type III fibril segments and then two classes with a potentially different but less-resolved structure were selected together for type IV with 77,667 segments. Type III morphology required just one more round of 3D classification to yield 9,831 ordered segments, which, after helical searches and CTF refinement, led to a final refined map at a resolution of 3.51 Å (0.143 FSC). The refined twist and twist were 358.99° and 4.83 Å respectively and the map was deposited with a sharpening B-factor of -74 Å^2^. For type IV morphologies, a map with resolved helical layer separation emerged from 5,955 segments after multiple further 3D classifications and refinements with helical searches. The final type IV map refined to a resolution of 3.8 Å with a helical twist and rise of 358.84° and 4.83 Å respectively and a sharpening B-factor of -77 Å^2^.

### Model building and refinement

A de novo model was built first for one layer of the type I fibril core in Coot^46^ as this form had the best resolution map. To avoid biasing the models, the multiple solved ex vivo structures were not used to guide model building and were only used for comparative purposes once the models had been built. Both Ramachandran and rotamer outliers were monitored and minimised during building in Coot. The final built layer was then repeated and rigid body fit to generate a model for 3 layers of the fibril core and this was used for real space refinement against the deposited map in Phenix v1.17.1^47^. NCS restraints were applied to prevent divergence of repeating chains in the layers, but the individual protomers within a layer were left to refine independently. A similar approach was then used for modelling the type III and IV structures with related backbone conformations initially docked in from the other polymorphs before being edited in Coot. The final real space refined models for each respective structure were assessed using MolProbity^48^ and deposited, with the final model statistics summarised in **Table 1**.

### Cellular seeding assays

HEK-293 cells were cultured in DMEM medium, supplemented with 10% FBS, 1 mM sodium pyruvate and non-essential amino acids (Gibco), under an atmosphere of 5% CO_2_ at 37°C. Cells were plated at 15.000 cells/well (for 24h seeding) or 10.000 cells/well (for 48h seeding) in 96-well PhenoPlates (PerkinElmer) that were coated with poly-L-lysine (Sigma) for 30 min. Cells were transfected using lipofectamine 3000 according to the manufacturer’s protocol. First, cells were transfected with 50 ng plasmid expressing tauRD-eYFP conjugates with a P301S mutation, w/o APR deletions. Following plasmid transfection (24h), cells were transfected further with rTau (24h incubation), peptide or AD-extracted aggregates (48h incubation). Just before transfection, samples were sonicated for 15 min (30 sec on, 30 sec off at 10°C) with a Bioruptor Pico (Diagenode). Then, 5 μL of sample, mixed with 0.2 ul of 3000 reagent, was added to a mixture of 4.5 μL of Opti-MEM medium (Gibco) with 0.3 μL Lipofectamine 3000. After a 15 min incubation at room temperature, 10 μL of mixture was added per well. Cell medium was replaced with 100% ice-cold methanol and plates were incubated on ice for 15 mins, then washed three times with PBS. Three individual plate preparations were performed per sample as independent experiments (n=3). High-content screening was performed at the VIB Screening Core/C-BIOS, using an Opera Phenix HCS (PerkinElmer) equipped with proper filter channels to track tau aggregation through the YFP channel (Ex:490-515nm, Em:525-580nm). Image storage (16 fields in 3 planes at a 40x magnification were acquired per well) and segmentation analysis was performed using the Columbus Plus digital platform (PerkinElmer).

### Preparation of recombinant tau seeds

Recombinant full-length tau (tau^2N4R^) was as described in previous protocols^49^. Lyophilised protein aliquots were freshly dissolved in 10mM HEPES, pH 7.4, 100mM NaCl at a final concentration of 10µM. Following filtration, using 0.2 µM PVDF filters, 5µM of heparin (Sigma) was added to the solution to induce aggregation. After 7 days of incubation at 37°C under shaking (700rpm), we generated seeds by breaking endpoint amyloid fibrils through successive sonication for 15min (30s on, 30s off) at 10°C, using a Bioruptor Pico sonication device (Diagenode).

### Extraction of tau aggregates

Ethical approval to access and work on the human tissue samples was given by the UZ Leuven ethical committee (Leuven/Belgium; File-No. S63759). An informed consent for autopsy and scientific use of autopsy tissue with clinical information was granted from all subjects involved (**Supplementary Table 1**). Following approval, brain tissue from autopsy cases was received from UZ/KU Leuven Biobank. Sarkosyl-insoluble material was extracted from cortex tissue of three individual patients with Alzheimer’s disease (AD1 – AD3), as described in previous work^8,50^. Tissue homogenisation was performed with a FastPrep (MP Biomedicals) in 10 volumes (w/v) cold buffer (10 mM Tris-HCl pH 7.4, 0.8 M NaCl, 1 mM EGTA and 10% sucrose) and a centrifugation step at 20,000 x g for 20 min at 4°C. Universal Nuclease (Pierce) was added to the supernatant, followed by a 30 min incubation at room temperature. Subsequently, the sample was brought to 1% Sarkosyl (Sigma) and incubated for 1h at room temperature while shaking (400 rpm), followed by centrifugation at 350,000 x g for 1h at 4°C. The pellet was washed once, resuspended in 50 mM Tris-HCl pH 7.4 (175 mg of starting material per 100 µl) and stored at -80°C.

## Data availability

The full atomic coordinates of the structures determined in this work will be available from the world wide Protein DataBase (wwPDB). All other data are available upon reasonable request.

## Acknowledgements

The authors gratefully acknowledge the Electron Microscopy Platform & Bio Imaging Core, Department of Neurosciences KU Leuven, VIB – KU Leuven Center for Brain & Disease Research for their support & assistance in this work. We also thank the VIB Screening Core/C-BIOS facility for excellent support with cellular screening. This work was supported by the Flanders institute for Biotechnology (VIB); KU Leuven; the Fund for Scientific Research Flanders (FWO, project grants G0C2818N, G0C0320N, and G053420N, FWO/Hercules Foundation equipment grants FWO AKUL/15/34 - G0H1716N, I011620N, and NextGenQBio - AH2016.133, and Postdoctoral Fellowships 12P0919N and 12P0922N to NL); the Stichting Alzheimer Onderzoek (SAO-FRA 2019/0015, SAO-FRA 2020/0009, and SAO-FRA 2020/0013). Neuropathological characterization of human brain samples was supported by FWO grants (G0F8516N and G065721N) to DRT.

## References

1. Kovacs, G.G. Tauopathies. Handb Clin Neurol 145, 355–368 (2017).

2. Goedert, M., Spillantini, M.G., Jakes, R., Rutherford, D. & Crowther, R.A. Multiple isoforms of human microtubule-associated protein tau: sequences and localization in neurofibrillary tangles of Alzheimer’s disease. Neuron 3, 519–26 (1989).

3. Jeganathan, S., von Bergen, M., Mandelkow, E.M. & Mandelkow, E. The natively unfolded character of tau and its aggregation to Alzheimer-like paired helical filaments. Biochemistry 47, 10526–39 (2008).

4. Gong, C.X. & Iqbal, K. Hyperphosphorylation of microtubule-associated protein tau: a promising therapeutic target for Alzheimer disease. Curr Med Chem 15, 2321–8 (2008).

5. Wesseling, H. et al. Tau PTM Profiles Identify Patient Heterogeneity and Stages of Alzheimer’s Disease. Cell 183, 1699–1713 e13 (2020).

6. Clavaguera, F. et al. Transmission and spreading of tauopathy in transgenic mouse brain. Nat Cell Biol 11, 909–13 (2009).

7. Frost, B., Jacks, R.L. & Diamond, M.I. Propagation of tau misfolding from the outside to the inside of a cell. J Biol Chem 284, 12845–52 (2009).

8. Louros, N. et al. Mapping the sequence specificity of heterotypic amyloid interactions enables the identification of aggregation modifiers. Nat Commun 13, 1351 (2022).

9. Abskharon, R. et al. Cryo-EM structure of RNA-induced tau fibrils reveals a small C-terminal core that may nucleate fibril formation. Proc Natl Acad Sci U S A 119, e2119952119 (2022).

10. Narasimhan, S. et al. Pathological Tau Strains from Human Brains Recapitulate the Diversity of Tauopathies in Nontransgenic Mouse Brain. J Neurosci 37, 11406–11423 (2017).

11. Scheres, S.H., Zhang, W., Falcon, B. & Goedert, M. Cryo-EM structures of tau filaments. Curr Opin Struct Biol 64, 17–25 (2020).

12. Shi, Y. et al. Structure-based classification of tauopathies. Nature 598, 359–363 (2021).

13. Arakhamia, T. et al. Posttranslational Modifications Mediate the Structural Diversity of Tauopathy Strains. Cell 180, 633–644 e12 (2020).

14. Wegmann, S., Medalsy, I.D., Mandelkow, E. & Muller, D.J. The fuzzy coat of pathological human Tau fibrils is a two-layered polyelectrolyte brush. Proc Natl Acad Sci U S A 110, E313–21 (2013).

15. Buee, L., Bussiere, T., Buee-Scherrer, V., Delacourte, A. & Hof, P.R. Tau protein isoforms, phosphorylation and role in neurodegenerative disorders. Brain Res Brain Res Rev 33, 95–130 (2000).

16. von Bergen, M. et al. Mutations of tau protein in frontotemporal dementia promote aggregation of paired helical filaments by enhancing local beta-structure. J Biol Chem 276, 48165–74 (2001).

17. von Bergen, M. et al. Assembly of τ protein into Alzheimer paired helical filaments depends on a local sequence motif (306VQIVYK311) forming β structure. Proceedings of the National Academy of Sciences 97, 5129–5134 (2000).

18. van der Kant, R., Louros, N., Schymkowitz, J. & Rousseau, F. Thermodynamic analysis of amyloid fibril structures reveals a common framework for stability in amyloid polymorphs. Structure (2022).

19. Falcon, B. et al. Structures of filaments from Pick’s disease reveal a novel tau protein fold. Nature 561, 137–140 (2018).

20. Fitzpatrick, A.W.P. et al. Cryo-EM structures of tau filaments from Alzheimer’s disease. Nature 547, 185–190 (2017).

21. Vaquer-Alicea, J., Diamond, M.I. & Joachimiak, L.A. Tau strains shape disease. Acta Neuropathol 142, 57–71 (2021).

22. Maurer-Stroh, S. et al. Exploring the sequence determinants of amyloid structure using position-specific scoring matrices. Nat Methods 7, 237–42 (2010).

23. Lovestam, S. et al. Assembly of recombinant tau into filaments identical to those of Alzheimer’s disease and chronic traumatic encephalopathy. Elife 11(2022).

24. Zhang, W. et al. Heparin-induced tau filaments are polymorphic and differ from those in Alzheimer’s and Pick’s diseases. Elife 8(2019).

25. Louros, N., van der Kant, R., Schymkowitz, J. & Rousseau, F. StAmP-DB: A platform for structures of polymorphic amyloid fibril cores. Bioinformatics (2022).

26. Schymkowitz, J. et al. The FoldX web server: an online force field. Nucleic Acids Res 33, W382–8 (2005).

27. Louros, N., Orlando, G., De Vleeschouwer, M., Rousseau, F. & Schymkowitz, J. Structure-based machine-guided mapping of amyloid sequence space reveals uncharted sequence clusters with higher solubilities. Nat Commun 11, 3314 (2020).

28. Arad, E., Green, H., Jelinek, R. & Rapaport, H. Revisiting thioflavin T (ThT) fluorescence as a marker of protein fibrillation – The prominent role of electrostatic interactions. Journal of Colloid and Interface Science 573, 87–95 (2020).

29. Ryder, B.D., Wydorski, P.M., Hou, Z. & Joachimiak, L.A. Chaperoning shape-shifting tau in disease. Trends in Biochemical Sciences 47, 301–313 (2022).

30. Chen, D. et al. Tau local structure shields an amyloid-forming motif and controls aggregation propensity. Nat Commun 10, 2493 (2019).

31. Arakhamia, T. et al. Posttranslational Modifications Mediate the Structural Diversity of Tauopathy Strains. Cell 184, 6207–6210 (2021).

32. Falcon, B. et al. Tau filaments from multiple cases of sporadic and inherited Alzheimer’s disease adopt a common fold. Acta Neuropathol 136, 699–708 (2018).

33. Zhang, W. et al. Novel tau filament fold in corticobasal degeneration. Nature 580, 283–287 (2020).

34. Louros, N., Schymkowitz, J. & Rousseau, F. Heterotypic amyloid interactions: Clues to polymorphic bias and selective cellular vulnerability? Curr Opin Struct Biol 72, 176–186 (2022).

35. Seidler, P.M. et al. CryoEM reveals how the small molecule EGCG binds to Alzheimer’s brain-derived tau fibrils and initiates fibril disaggregation. bioRxiv, 2020.05.29.124537 (2020).

36. Li, X. et al. The distinct structural preferences of tau protein repeat domains. Chem Commun (Camb) 54, 5700–5703 (2018).

37. Haj-Yahya, M. et al. Site-Specific Hyperphosphorylation Inhibits, Rather than Promotes, Tau Fibrillization, Seeding Capacity, and Its Microtubule Binding.

38. Leonard, C., Phillips, C. & McCarty, J. Insight Into Seeded Tau Fibril Growth From Molecular Dynamics Simulation of the Alzheimer’s Disease Protofibril Core. Frontiers in Molecular Biosciences 8(2021).

39. Jucker, M. & Walker, L.C. Propagation and spread of pathogenic protein assemblies in neurodegenerative diseases. Nat Neurosci 21, 1341–1349 (2018).

40. von Bergen, M. et al. Assembly of tau protein into Alzheimer paired helical filaments depends on a local sequence motif ((306)VQIVYK(311)) forming beta structure. Proc Natl Acad Sci U S A 97, 5129–34 (2000).

41. Ambadipudi, S., Reddy, J.G., Biernat, J., Mandelkow, E. & Zweckstetter, M. Residue-specific identification of phase separation hot spots of Alzheimer’s-related protein tau. Chem Sci 10, 6503–6507 (2019).

42. Savitzky, A. & Golay, M.J.E. Smoothing and Differentiation of Data by Simplified Least Squares Procedures. Analytical Chemistry 36, 1627–1639 (1964).

43. Zheng, S.Q. et al. MotionCor2: anisotropic correction of beam-induced motion for improved cryo-electron microscopy. Nat Methods 14, 331–332 (2017).

44. Zhang, K. Gctf: Real-time CTF determination and correction. J Struct Biol 193, 1–12 (2016).

45. Wagner, T. & Raunser, S. The evolution of SPHIRE-crYOLO particle picking and its application in automated cryo-EM processing workflows. Communications Biology 3, 61 (2020).

46. Emsley, P., Lohkamp, B., Scott, W.G. & Cowtan, K. Features and development of Coot. Acta Crystallogr D Biol Crystallogr 66, 486–501 (2010).

47. Adams, P.D. et al. PHENIX: a comprehensive Python-based system for macromolecular structure solution. Acta Crystallogr D Biol Crystallogr 66, 213–21 (2010).

48. Williams, C.J. et al. MolProbity: More and better reference data for improved all-atom structure validation. Protein Sci 27, 293–315 (2018).

49. Mirbaha, H. et al. Inert and seed-competent tau monomers suggest structural origins of aggregation. Elife 7(2018).

50. Goedert, M., Spillantini, M.G., Cairns, N.J. & Crowther, R.A. Tau proteins of Alzheimer paired helical filaments: abnormal phosphorylation of all six brain isoforms. Neuron 8, 159–68 (1992).

