## Supplementary information for "Tau amyloid polymorphism is shaped by local structural propensities of its protein sequence"

This PDF file includes:
Supplementary Figures 1-2

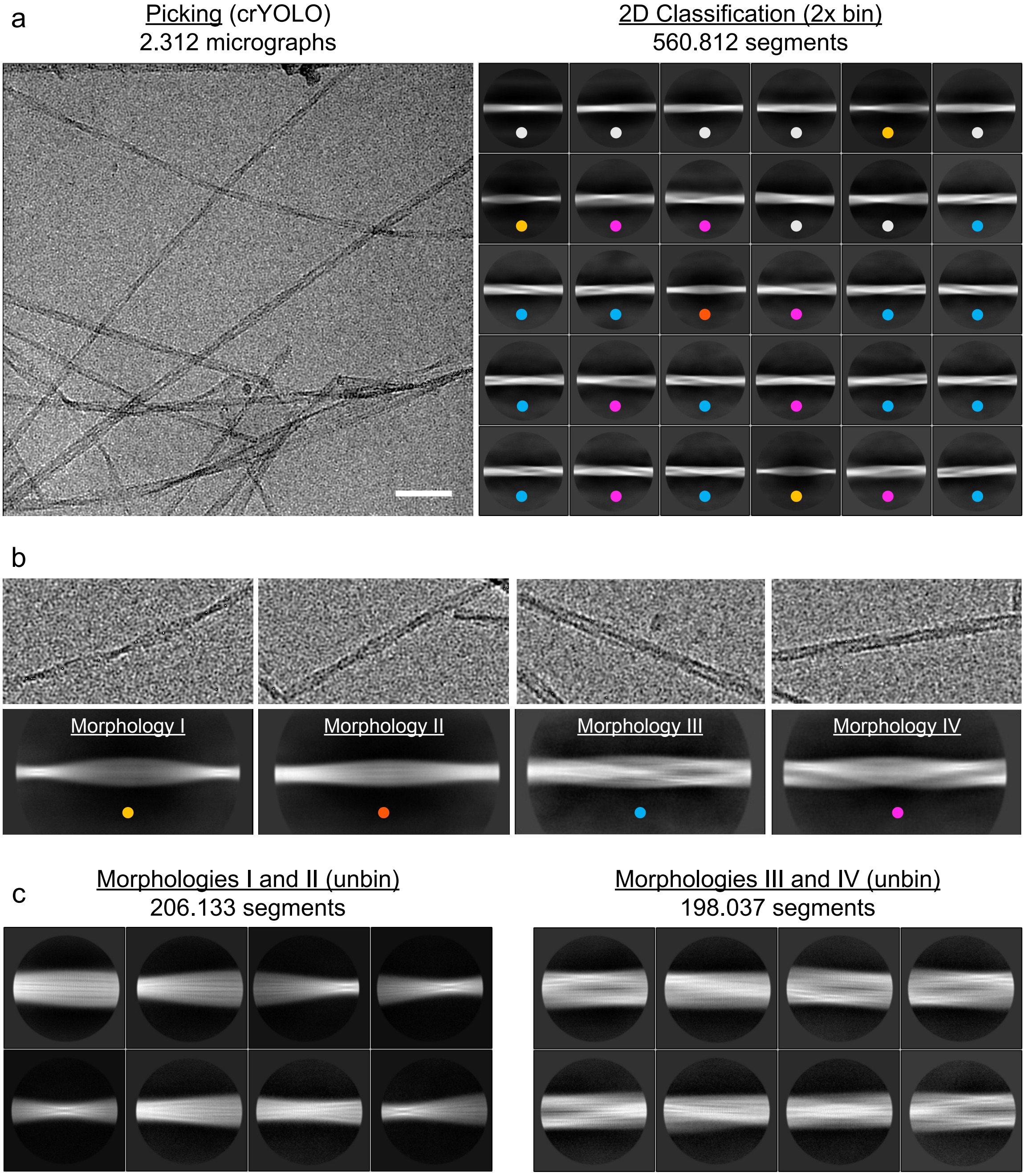


**Supplementary Figure 1. Initial analysis of the Tau3 cryo-EM dataset revealed multiple distinct fibril polymorphs.** (a) An example micrograph from the cryo-EM dataset and the subsequent most populated 2D class averages from all of the extracted fibril segments highlight the presence of multiple distinct fibril polymorphs. Consistent forms that could be identified from the 2D class averages are colour-coded to later images whereas ambiguous classes are labelled with grey circles. (b) At this stage, four major polymorphs were identified with representative 2x binned class averages below close-up sections of a corresponding fibril from the raw micrographs. (c) The data was split into two subsets based on the fibril morphology apparent in 2D class averages and example higher resolution 2D class averages are displayed from unbinned fibril segments.

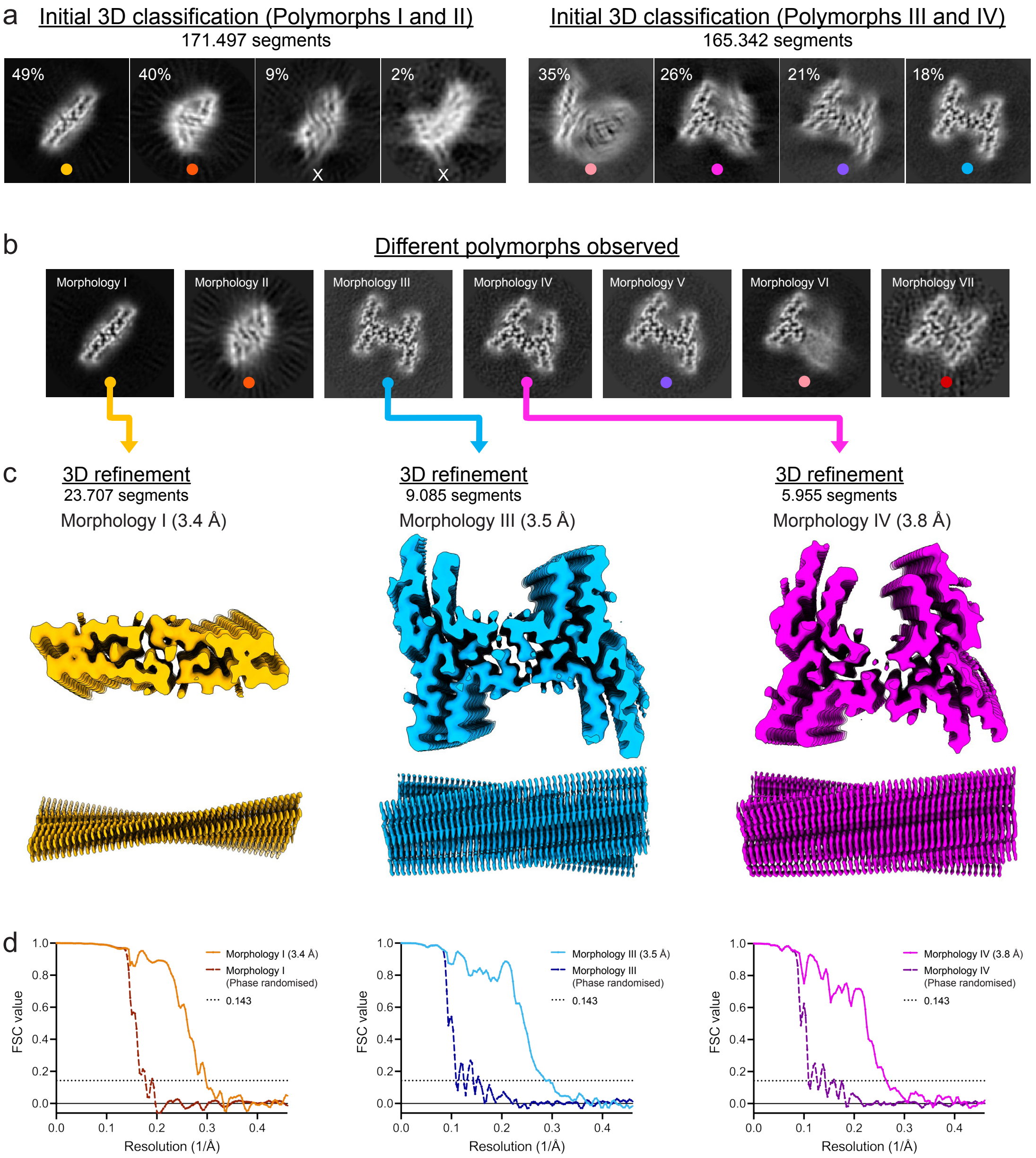


**Supplementary Figure 2. Continued processing of the Tau$_{350\text{-}362}$ cryo-EM dataset led to the solution of multiple related fibril structures.** (a) The first 3D classification round for each of the two segment subsets started to show different fibril core structures within the data. (b) Multiple rounds of 3D classification and helical parameter optimisations were subsequently required before the peptide backbone of most of the different polymorphs could be resolved. The best resulting classes for each form are shown as sums of the central 6x slices through each respective map. (c) From the seven polymorphs identified, three led to high resolution maps where the helical β-strand layers could be completely resolved, deposited as EMDB XXXX (Morphology I), EMDB XXXX (Morphology III) and EMDB XXXX (Morphology IV) respectively. (d) The FSC plots with the corrected FSC and phase randomised values of each of the solved forms are shown.

**Supplementary Table 1.** Basic information of the patients and disease stage. An informed consent for autopsy and scientific use of autopsy tissue with clinical information was granted from all subjects involved.

| Sample ID | Age (years) | Sex | Neuropathological diagnosis | Braak NFT stage | Aβ phase | CERAD score | NIA-AA degree of AD pathology | Post Mortem Interval/hours |
|---|---|---|---|---|---|---|---|---|
| AD1 | 71 | Female | AD | VI | 5 | 2 | high | 24 |
| AD2 | 87 | Male | AD | VI | 5 | 2 | high | 12 |
| AD3 | 71 | Male | AD | V | 5 | 3 | high | 12 |
